# Cul4-Ddb1 ubiquitin ligases facilitate DNA replication-coupled sister chromatid cohesion through regulation of cohesin acetyltransferase Esco2

**DOI:** 10.1101/414888

**Authors:** Haitao Sun, Jiaxin Zhang, Jingjing Zhang, Zhen Li, Qinhong Cao, Huiqiang Lou

**Author notes:** To whom correspondence should be addressed: Qinhong Cao; (QC); and Huiqiang Lou; (HL).

## Abstract

Cohesin acetyltransferases Esco1 and Esco2 play a vital role in establishing sister chromatid cohesion. How Esco1 and Esco2 are controlled to achieve this in a DNA replication-coupled manner remains unclear in higher eukaryotes. Here we show that <u>Cul4</u>-<u>R</u>ING ligases (CRL4s) play a critical role in sister chromatid cohesion in human cells. Depletion of Cul4A, Cul4B or Ddb1 subunits substantially reduces normal cohesion efficiency. We also show that Mms22L, a vertebrate ortholog of yeast Mms22, is one of <u>Ddb1</u> and <u>Cul</u>4-<u>a</u>ssociated <u>f</u>actors (DCAFs) involved in cohesion. Several lines of evidence suggest a selective interaction of CRL4s with Esco2, but not Esco1. Depletion of either CRL4s or Esco2 causes a defect in Smc3 acetylation which can be rescued by HDAC8 inhibition. More importantly, both CRL4s and PCNA act as mediators for efficiently stabilizing Esco2 on chromatin and catalyzing Smc3 acetylation. Taken together, we propose an evolutionarily conserved mechanism in which CRL4s and PCNA regulate Esco2-dependent establishment of sister chromatid cohesion.

**Author summary:** We identified human Mms22L as a substrate specific adaptor of Cul4-Ddb1 E3 ubiquitin ligase. Downregulation of Cul4A, Cul4B or Ddb1 subunit causes reduction of acetylated Smc3, via interaction with Esco2 acetyltransferase, and then impairs sister chromatid cohesion in 293T cells. We found functional complementation between Cul4-Ddb1-Mms22L E3 ligase and Esco2 in Smc3 acetylation and sister chromatid cohesion. Interestingly, both Cul4-Ddb1 E3 ubiquitin ligase and PCNA contribute to Esco2 mediated Smc3 acetylation. To summarise, we demonstrated an evolutionarily conserved mechanism in which Cul4-Ddb1 E3 ubiquitin ligases and PCNA regulate Esco2-dependent establishment of sister chromatid cohesion.

## Introduction

Faithful inheritance of the genetic information requires precise chromatin replication and separation of sister chromatids into two daughter cells. To ensure accurate chromosome segregation in eukaryotic cells, a pair of sister chromatids should be aligned properly and held together by a cohesin complex from S phase to anaphase [1–6]. The cohesin complex is a four-subunit ring conserved from yeast to human. In human mitotic cells, cohesin is composed of Smc1, Smc3, Rad21 (Scc1/Mcd1 in yeast) and SA1 or SA2 (Scc3 in yeast) [2, 7–10].

Cohesin is widely believed to have distinct statuses according to its association with chromatin during the cell cycle. In G_1_ phase, it is loaded loosely onto chromatin (i.e., non-cohesive status) [11]. As cells proceed into S phase, cohesin binds more tightly to hold sister chromatids together (i.e. cohesive status), and this transition is called the establishment of sister chromatid cohesion [5, 12]. Although the structural bases of this transition remain enigmatic, it has been shown in yeast (*Saccharomyces cerevisiae*) to depend on an essential cohesin acetyltransferase, Eco1 [13–15]. Eco1 triggers cohesion establishment during S phase through counteracting the opposing activity of Rad61 (WAPL in human) [16]. The essential substrate of Eco1 has been demonstrated to be Smc3 [13].

Cohesion is established in a DNA replication-coupled manner [12, 17, 18]. To achieve this, the activity of Eco1 is controlled concomitantly with DNA replication by two independent mechanisms. First, Eco1 contains a canonical PIP (PCNA interaction peptide) box, which mediates its interaction with PCNA, the multivalent-platform of DNA replisome [19]. Second, a member of the cullin-RING E3 ligases (CRLs) Rtt101-Mms1, associates with the replication fork and facilitates Smc3 acetylation through direct association between Eco1 and the substrate receptor component of the ligase, Mms22 [20].

CRLs constitute the largest ubiquitin ligase family in eukaryotes. They are modular assemblies consisting of a Cullin scaffold in complex with an adapter and distinct ligase substrate receptors, giving rise to many combinatorial possibilities. There are three cullins in budding yeast (Cul1, 3 and 8) and six in human (Cul1, 2, 3, 4A, 4B and 5) [21]. Cul8, also known as Rtt101, is unique to budding yeast, but shows low sequence similarity with Cul4. Nevertheless, the Rtt101 adaptor Mms1 is highly homologous to human Ddb1 adapter of Cul4, and Rtt101-Mms1 performs similar functions to Cul4-Ddb1 ligases in other organisms [22]. The human genome encodes two Cul4 paralogs, Cul4A and Cul4B, sharing 80% sequence identity aside from Cul4B having an extended N-terminus containing a nuclear localization signal (NLS) [23]. Both Cul4A and Cul4B use Ddb1 as an adaptor and DCAFs (<u>Ddb1</u> and <u>Cul4</u>-<u>a</u>ssociated <u>f</u>actors) as substrate receptors to recognize a large number of substrate proteins [24–26]. CRL4s play recognized roles in DNA repair, replication and chromatin modifications through ubiquitylation and/or mediating protein-protein interactions [27, 28].

Mammalian cells have two Eco1 orthologs, Esco1 and Esco2 [29]. Both of these have been shown to acetylate Smc3 at two evolutionarily conserved lysine residues (K105K106) [15, 30, 31]. Interestingly, Esco1 acetylates Smc3 through a mechanism distinct from that of Esco2 [32]. Nevertheless, how the activities of Esco1 and Esco2 are controlled to establish replication-coupled sister chromatid cohesion in vertebrates has not been delineated.

In this study, we report that Cul4-Ddb1 E3 ligases function in establishing sister chromatid cohesion in human cells. Depletion of Cul4A, Cul4B or Ddb1 results in precocious sister chromatid separation. We show that Mms22L (Mms22-like), the human ortholog of yeast Mms22, the substrate receptor of Rtt101-Mms1, interacts with Ddb1. Interestingly, Esco2, not Esco1, co-immunoprecipitates with all of the subunits of CRL4^Mms22L^ ligase. Dosage suppression experiments reveal that CRL4s and Esco2 are able to compensate each other in Smc3 acetylation and thereby sister chromatid cohesion in 293T cells. Through introducing interaction defective mutations, we find that Esco2 acetylates Smc3 dependent on interactions with both Cul4-Ddb1 ligases and PCNA. These data suggest that Cul4-Ddb1 ligases and PCNA contribute together to connect Esco2-dependent cohesion establishment with the replication process in human.

## Results

### Cul4-Ddb1 and Mms22L are required for efficient sister chromatid cohesion in human cells

Recently, we showed that fork-associated Rtt101-Mms1 ubiquitin ligases function in linking the establishment of sister chromatid cohesion with DNA replication in yeast [20]. We asked whether Cul4-Ddb1, the putative functional homolog of Rtt101-Mms1 in human cells, participate in sister chromatid cohesion as well. To test this, we depleted Cul4A, Cul4B or Ddb1 from 293T cells using small interfering RNA (siRNA) and measured sister chromatid cohesion. Cultured cells were harvested by trypsinization to enrich for cells in mitosis. Chromosome spreads were stained with Giemsa and the morphology of the mitotic cells was analyzed (Fig 1A). We did not synchronize cells in metaphase with nocodazole since vertebrate cohesins are removed from chromosome arms in prophase allowing only cohesion of centromeres to be monitored [33]. We, however, wished to monitor cohesion not only at centromeres but also at telomeres and chromosome arms, where Rtt101-Mms1 have been shown to be required for cohesion establishment [20] (Fig 1A). In our experiments, “normal cohesion” defines the state in which both centromere and chromosome arms are closely tethered each other (i, Fig 1A), whereas arm open (ii), partially separated but still paired (also called “railroad”, iii), unpaired (iv) or completely separated (v) chromatids indicate various extents of cohesion impairment. Under our experimental conditions, most chromatids in a single cell display similar morphology. We calculated the cohesion percentages as the proportion of “normal cohesion” cells (i) among total mitotic cells, where indicated. Alternatively, the percentage of cells bearing separated centromeres (iii, iv and v, Fig 1A) was used as an indicator as severe cohesion deficiency (S1 Fig). Depletion of either Cul4A or Cul4B reduced the “normal cohesion” from ∼80% to ∼40% (Figs 1B and 1C). The specificity of RNA interference (RNAi) was verified through complementation by over-expressing the respective proteins carrying a Flag tag. This indicates that Cul4A and Cul4B may play at least partially non-redundant roles in sister chromatid cohesion. Similar results were observed for cells devoid of Ddb1, whereas the cell cycle progression was not significantly affected (Figs 1D and S1A-B). These results indicate that Cul4A, Cul4B and Ddb1, like their homologs in yeast, are required for efficient cohesion in human cells.

**Fig 1.**
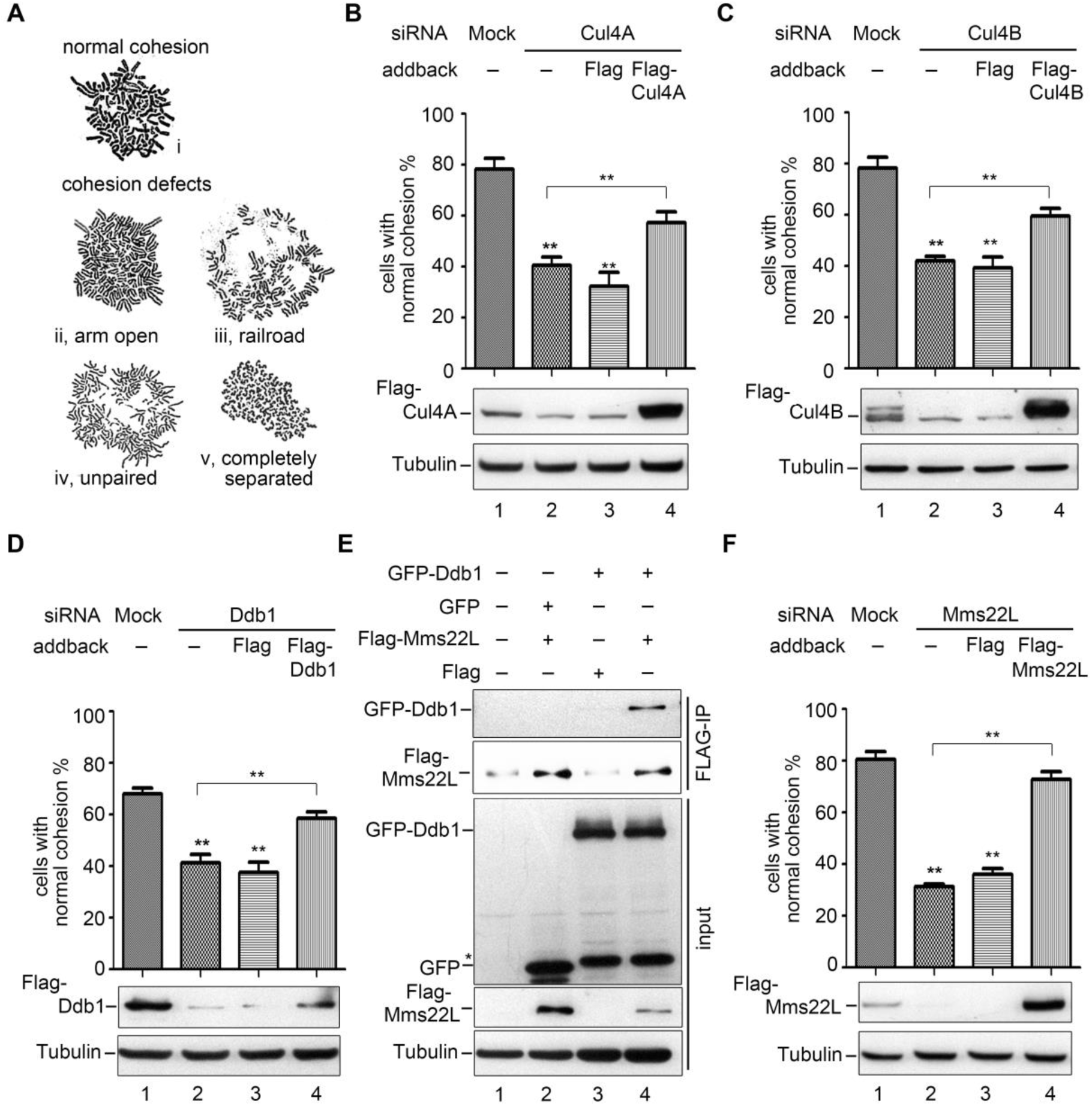
Knockdown of Cul4-Ddb1-Mms22L causes severe cohesion defects. (A) Representative morphologies of human chromosome spreads stained with Giemsa. Closed sister chromatids (i) indicates normal cohesion, while loose sister chromatids (arm opened, loosely paired, unpaired and completely separated, ii-v) indicate different extents of cohesion defects. We have only rarely observed that chromosomes within one cell display different morphologies. In order to reflect the physiological status of chromosome morphologies, cells were not synchronized. At least 200 mitotic cells were enriched and harvested via trypsin digestion for each experiment. The percentage of cells with closed sister chromatids among the total mitotic cells (i.e., normal cohesion %) was quantified from at least three independent experiments. (B-D) Cohesion defects caused by depletion of Cul4A (B), Cul4B (C) and Ddb1 (D). 293T cells were transfected with siRNAs specific to Cul4A, Cul4B and Ddb1 for 48 h. A plasmid expressing the indicated Flag-tagged protein was transferred into the cells after 6 h for the complementation assay. The trysinized cells were fixed with methanol and acetic acid (3:1) for three times and then stained with Giemsa. More than 200 mitotic cells per RNAi experiment were scored; the results of at least three independent biological experiments were summarized in the histogram. The cohesion percentage of each RNAi sample was compared with that of mock or add-back using student’s *t*-test, **P<0.01. The efficiency of siRNA and complementation of Cul4A, Cul4B, Ddb1 or Mms22L was detected via immunoblots against the indicated antibodies. (E) Ddb1 co-precipitates with Mms22L. *Flag, Flag-Mms22L* and *GFP, GFP-Ddb1* plasmids were transferred into 293T cells. After IP experiments, Mms22L and Ddb1 were detected with antibodies against Flag and GFP, respectively. Tubulin was probed as a loading control. (F)Mms22L depletion leads to compromised cohesion as well. Quantification of the cohesion percentage was performed as described above. See Supporting S1 Fig for the *CEN* cohesion defect results.

Mms22 is one of the substrate adaptors of Rtt101-Mms1 in yeast. Mms22L, a putative human ortholog of Mms22, functions together with Cul4-Ddb1 in replication-coupled nucleosome assembly [34]. However, it remains unknown whether it is a DCAF to date. To test this, we next co-expressed GFP-Ddb1 and Flag-Mms22L in 293T cells. Flag-Mms22L was immunoprecipitated by anti-Flag antibodies from whole cell extracts. As shown in Fig 1E, considerable amounts of Ddb1 co-precipitated with Flag-Mms22L, arguing that Mms22L interacts with Ddb1 and is likely a new DCAF of CRL4 ligases in human. Interestingly, Mms22L depletion resulted in significant cohesion defects at both chromosome arms and centromeres, reminiscent of depletion of other CRL4 subunits Cul4A, Cul4B or Ddb1 (Figs 1F and S1C). Taken together, these data suggest that CRL4^Mms22L^ ligases participate in sister chromatid cohesion in human cells.

### CRL4^Mms22L^ ligases selectively interact with Esco2

To answer how CRL4s affect sister chromatid cohesion, we tested whether CRL4 subunits interact with the key cohesin acetyltransferases Esco1 or Esco2. We first observed the cellular distribution of Esco1, Esco2, Cul4A, Cul4B or Ddb1 by immunofluorescence. RFP-labelled Esco1 or Esco2 and GFP-tagged Cul4A, Cul4B or Ddb1 were introduced into 293T cells. In agreement with previous observations [23, 35], Cul4B localized to nucleus, whereas Cul4A and Ddb1 distributed throughout the whole cell. Esco1 and Esco2 mainly distributed within nucleus (Figs 2A and 2B), as reported previously by other groups [29, 36].

**Fig 2.**
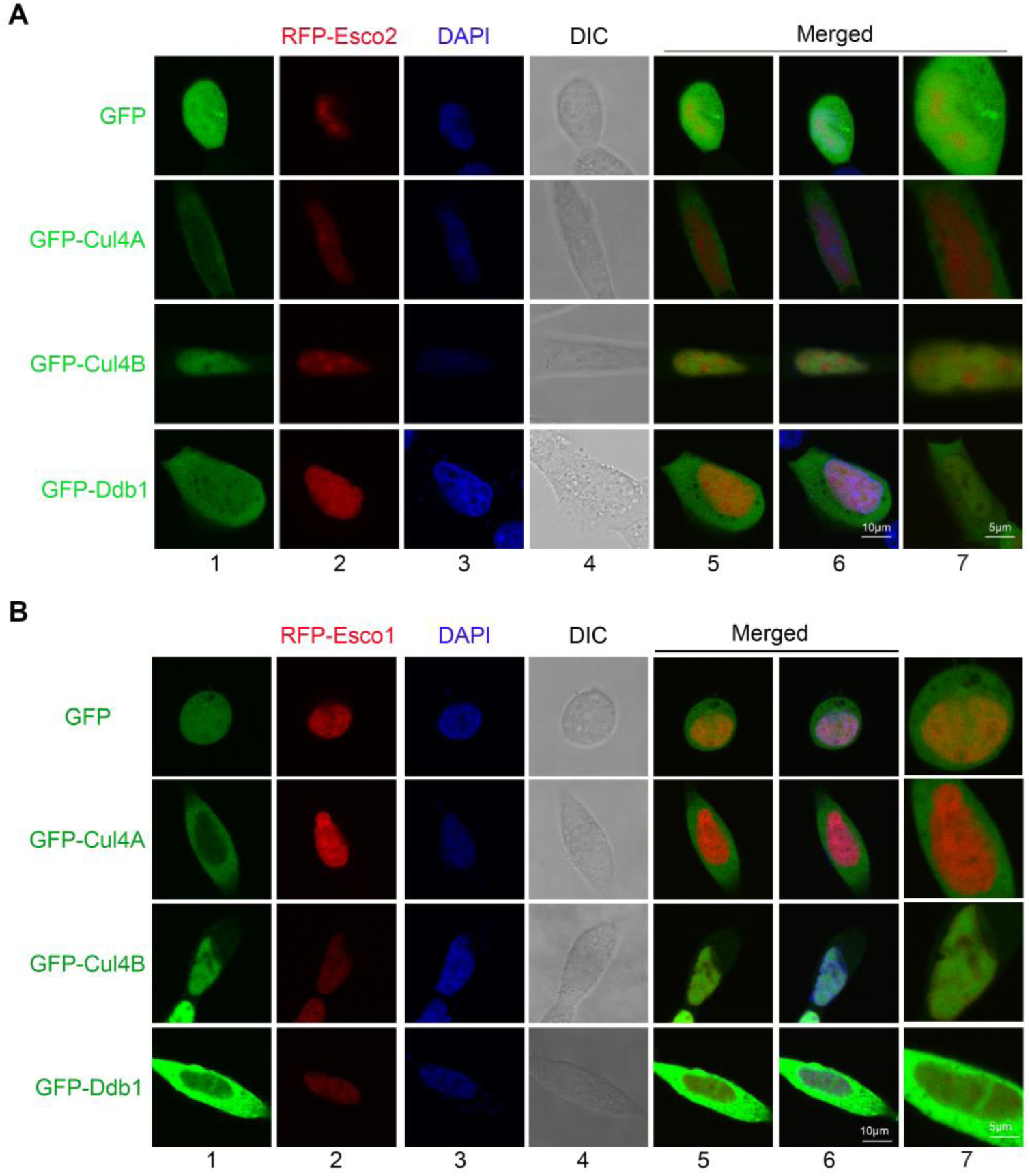
Esco2 co-localizes with Cul4 and Ddb1. (A) Localization of Esco2, Cul4A, Cul4B and Ddb1. 293T cells were co-transferred with *RFP-Esco2* plasmids and *GFP, GFP-Cul4A, GFP-Cul4B*, or *GFP-Ddb1* plasmids. After 24 h, nuclei were stained with DAPI. Pictures were captured with a laser-confocal microscope. RFP and GFP images were merged with (lane 6) or without DAPI (lane 5). (B) Esco1 does not co-localize with CRL4s. 293T cells were co-transferred with *RFP-Esco1* plasmids and *GFP, GFP-Cul4A, GFP-Cul4B*, or *GFP-Ddb1* plasmids. Fluorescence microscopy was conducted as described above.

Meanwhile, in order to obtain insight into how Esco2 is regulated, we searched for its interaction partners using affinity purification coupled mass spectrometry (AP-MS). To this end, Esco2 carrying both His6 and 5Flag tags was over-expressed in 293T cells and subjected to tandem affinity purification. Interestingly, Ddb1, together with many histone subunits and chaperones (e.g., HP1), was repeatedly detected among the co-purified proteins with Esco2-HF (Fig 3A). We then performed immunoprecipitations to corroborate the interaction through ectopically expressing Flag tagged subunit of CRL4s in 293T cells. Consistently, Esco2 clearly co-precipitated with not only Ddb1 (Fig 3B) but also other CRL4 subunits (Figs 3C and S2). On the contrary, virtually no Esco1 was detectable in the precipitates of any CRL4 subunits in all experiments carried out in parallel with Esco2 (Figs 3B-C and S2). These data indicate that CRL4 ligases might have a preferential association with Esco2.

**Fig 3.**
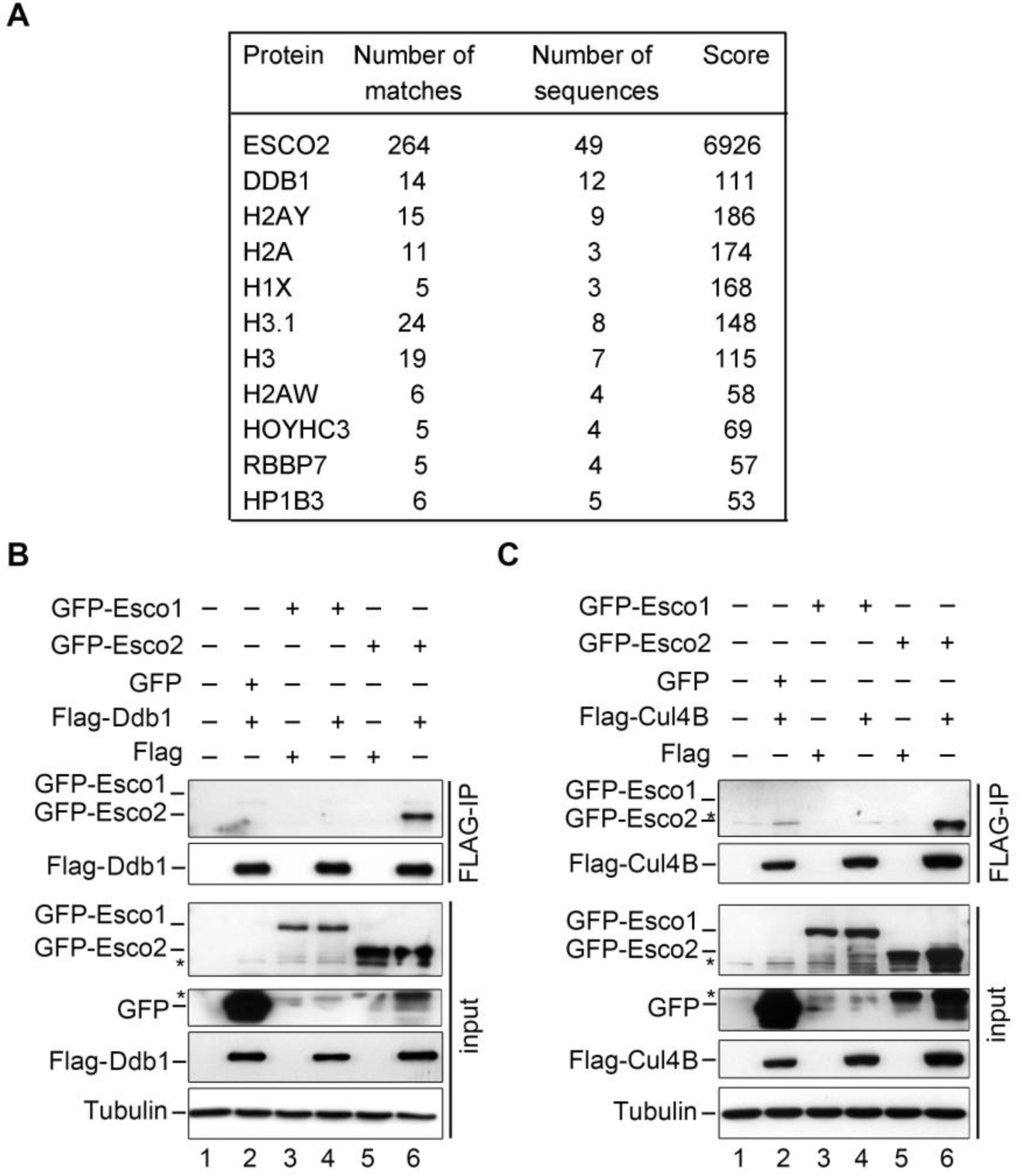
Preferential interaction between CRL4s^Mms22L^ and Esco2. (A) Ddb1 is repeatedly co-purified with Esco2. 293T cells were co-transferred with *Esco2-HF* plasmid. After tandem affinity purification, proteins in the final elution were analyzed by MS. Cells expressing Esco1-HF were conducted as a control. (B, C) Esco2, but not Esco1, co-immunoprecipitates with Cul4A-Cul4B-Ddb1-Mms22L. *GFP, GFP-Esco1, GFP-Esco2* and *Flag-Ddb1* (B) or *Flag-Cul4B* (C) were co-expressed in 293T cells. Flag-IP experiments were performed as described in Fig 1E. The asterisks indicate non-specific reacting bands. See Supporting S2 Figs for the data of *Flag-Cul4A* and *Flag-Mms22L* immunoprecipitation experiments.

### Esco2 functions in a CRL4^Mms22L^-dependent manner

Given the possible interaction between CRL4s and Esco2, we asked whether lack of Ddb1-Mms22L can be compensated by over-expressing Esco2. To test this, we ectopically expressed Esco2 in a Ddb1 (Fig 4A) or Mms22L depleted background (Fig 4B). Both mild and severe cohesion defects in either Ddb1 or Mms22L-depleted 293T cells were markedly rescued by over-expression of Esco2 (Figs 4A-B, S3A-B, lane 6), indicating a potent functional interaction between Ddb1^Mms22L^ and Esco2 as well as the physical interaction documented in Fig 3. We had previously isolated a separation-of-function mutant in yeast, *eco1-LG* (L61DG63D), which shows a dramatically compromised interaction with Mms22 [20]. Interestingly, these two residues are highly conserved in Esco2 (L415G417), but not in Esco1 (S3C Fig), which correlates well with their different abilities to interact with CRL4s^Mms22L^. Over-expression of *Esco2*-LG (*Esco2*-L415DG417D) mutant suppressed to a lesser extent than wild-type (WT) *Esco2* (Figs 4A-B, S3A-B, compare lane 7 to 6), indicating that the role of Esco2 is at least partially dependent on its interaction with Ddb1^Mms22L^. In order to further address the contribution of the interaction of Esco2 and CRL4s in sister chromatid cohesion, we tested the dosage suppression effects in an Esco2-depleted background. In comparison to WT Esco2, expression of the interaction defective mutant *Esco2*-LG hardly displayed suppression (Figs 4C and S3D, compare lane 6 to 5). This result reinforces the argument that the interaction between Esco2 and CRL4^Mms22L^ is important for the role of Esco2 in cohesion establishment. To further support this, Cul4, Ddb1 and Mms22L are dosage suppressors of Esco2 knockdown mutant as well (Figs 4C and S3D, lanes 7-10). These results implicate that Ddb1^Mms22L^ might serve as a critical positive regulator of the cohesion function of Esco2.

**Fig 4.**
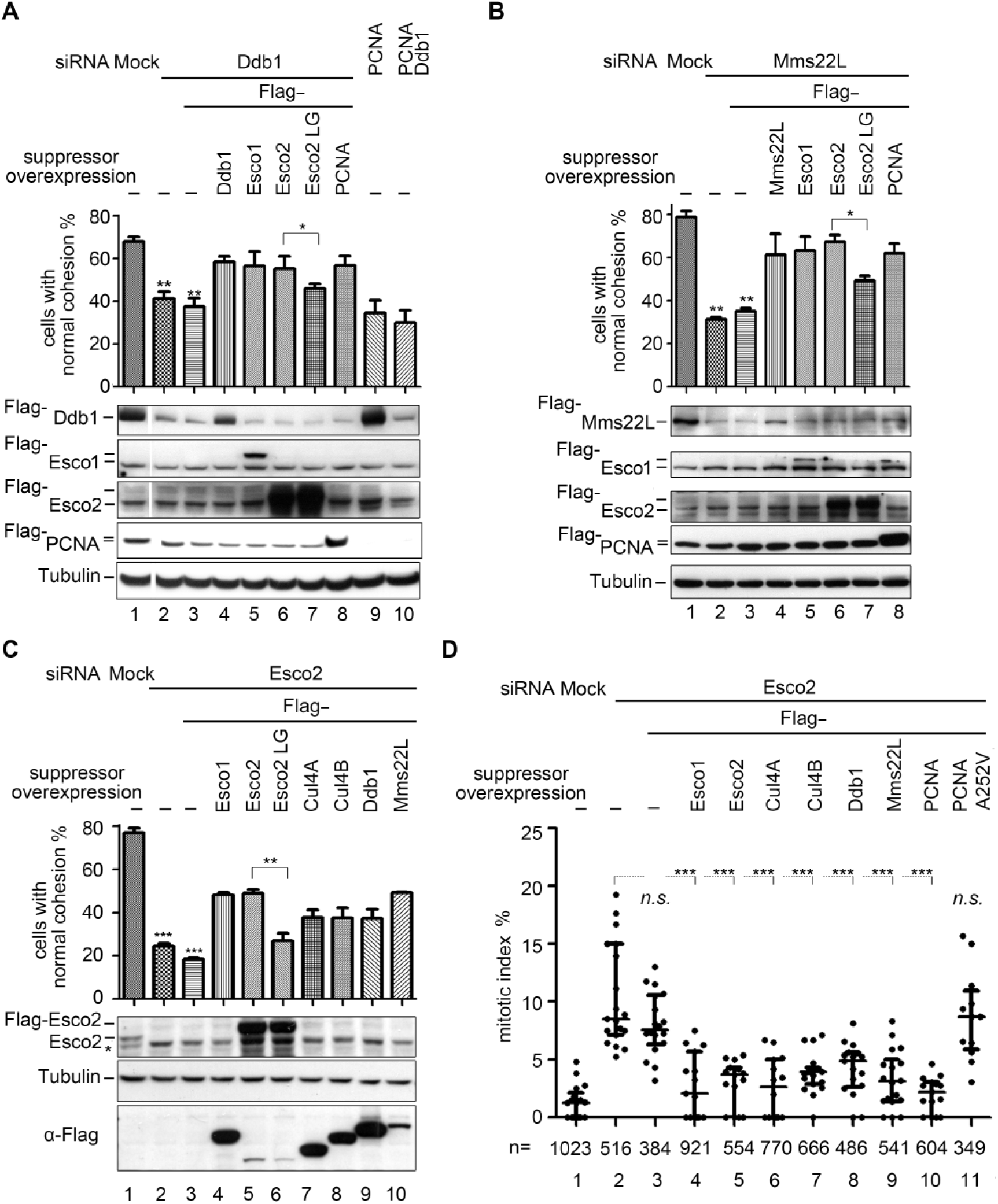
Functional interaction between CRL4s^Mms22L^ and Esco2. (A) Over-expression of Esco1, Esco2 or PCNA partially suppresses a cohesion defect in Ddb1 knockdown cells, while over-expression of *Esco2-LG* mutant does not. Immunoblots of Ddb1, Esco1, Esco2, PCNA and Tubulin were shown below each column. The statistical significance was calculated via student’s *t*-test, ** P<0.01; * P<0.05. The results of at least three independent biological experiments were summarized in the histogram. See Supporting S3A Fig for the corresponding *CEN* cohesion defect results. (B) Esco1, Esco2 or PCNA over-expression partially compensates a SCC defect caused by Mms22L knockdown, while over-expression of *Esco2-LG* mutant does not. Immunoblots of Mms22L, Esco1, Esco2, PCNA and Tubulin are shown below each column. The statistical significance was calculated via student’s *t*-test, ** P<0.01; * P<0.05. The results of at least three independent biological experiments were summarized in the histogram. See Supporting S3B Fig for the corresponding *CEN* cohesion defect results. (C) Over-expression of Cul4A, Cul4B, Ddb1, or Mms22L partially suppresses the SCC defect caused by Esco2 knockdown, while over-expression of an E3-interaction defective mutant, or *Esco2*-LG (L415DG417D) mutant has no effect. Esco2 was knocked down by siRNA in 293T cells, then Flag-tagged Esco1, Esco2, *Esco2*-LG, Cul4A, Cul4B, Ddb1, or Mms22L were overexpressed after siRNA delivery for 6 h. The percentage of cells in cohesion was determined as in Fig 1. Immunoblots of Esco2, Flag and Tubulin from each RNAi experiment are shown below the corresponding column. The statistical significance was calculated via student’s *t*-test, *** P<0.001; ** P<0.01. The results of at least three independent biological experiments were summarized in the histogram. See Supporting S3D Fig for the corresponding *CEN* cohesion defect results. See Supporting S3E Fig for the results of PCNA and PCNA-A252V alleles. (D) Over-expression of Cul4A, Cul4B, Ddb1, Mms22L or PCNA suppresses the M phase arrest caused by Esco2 knockdown. The portion of cells in the M phase among total cells was counted after Hoechst 33342 staining. The statistical significance was calculated via student’s *t*-test, *** P<0.001, n: the total cell number counted. The results of at least three independent biological experiments were summarized.

Because defects in sister chromatid cohesion often activate the spindle checkpoint and result in the G_2_/M arrest of the cell cycle, we then examined the proportion of M-phase cells (mitotic index). The mitotic index was very low for untreated 293T cells, but increased to an average about 12% when Esco2 was depleted (Fig 4D, column 2) consistent with the observations from another group [37]. The G_2_/M arrest induced by Esco2 knockdown was dramatically alleviated via over-expression of Cul4A or Cul4B (Fig 4D, columns 6 and 7), Ddb1 (column 8) or Mms22L (column 9). Taken together, these data demonstrate that the interaction between CRL4^Mms22L^ and Esco2 is important for Esco2 function in sister chromatid cohesion and thereby mitotic progression.

Besides interaction with CRL4s, the activity of Eco1 is also linked with replication forks through association with PCNA in yeast [19]. This notion was corroborated because the cohesion defects (S3E Fig) and mitotic arrest (Fig 4D, compare lanes 10 and 11) in Esco2-depleted 293T cells were significantly rescued by over-expression of WT PCNA, but not by an Esco-interaction defective mutant *PCNA*-A252V. Together, these data suggest that both CRL4^Mms22L^ and PCNA mediated interactions are critical for the Esco2-dependent establishment of sister chromatid cohesion.

### Cul4-Ddb1 ligases participate in sister chromatid cohesion by promoting Esco2-mediated Smc3 acetylation

Given that the essential role of Eco1/Esco lies in catalyzing Smc3 acetylation during cohesion establishment [13, 38], we next examined whether the dosage suppression effects observed above are due to facilitating Smc3 acetylation. For this purpose, Smc3 acetylation was measured in 293T cell lysates via immunoblots with an antibody that specifically recognizes Smc3K105ac/K106ac. S4A Fig demonstrates that the antibody recognizes an amount of Smc3ac proportional to the input protein concentrations. Esco2-depleted cells displayed substantially reduced Smc3 acetylation (S4B Fig, compare lanes 1, 2 and 7), which was partially restored through ectopic expression of Ddb1 or Mms22L (S4B Fig, compare lanes 1, 5 and 6). Over-expression of either Cul4A or Cul4B was also capable of stimulating Smc3 acetylation to a similar extent (S4B Fig, lanes 3 and 4). These results suggest that the compensation of cohesion defects in Esco2-depleted cells by Cul4, Ddb1, or Mms22L over-expression may be achieved through enhancing Smc3 acetylation.

Next, we determined whether CRL4s directly participate in regulating Smc3 acetylation. Depletion of each subunit of CRL4s led to moderately compromised Smc3 acetylation (Figs 5A and S4C-S4E), indicating that CRL4s^Mms22L^ are required for efficient Esco2-dependent Smc3 acetylation. Meanwhile, the protein levels of both Esco enzymes were not significantly affected (Fig 5A, descending panels 3 and 4), suggesting that Cul4-Ddb1-Mms22L unlikely regulate the expression and/or protein turnover of Esco1 and Esco2.

**Fig 5.**
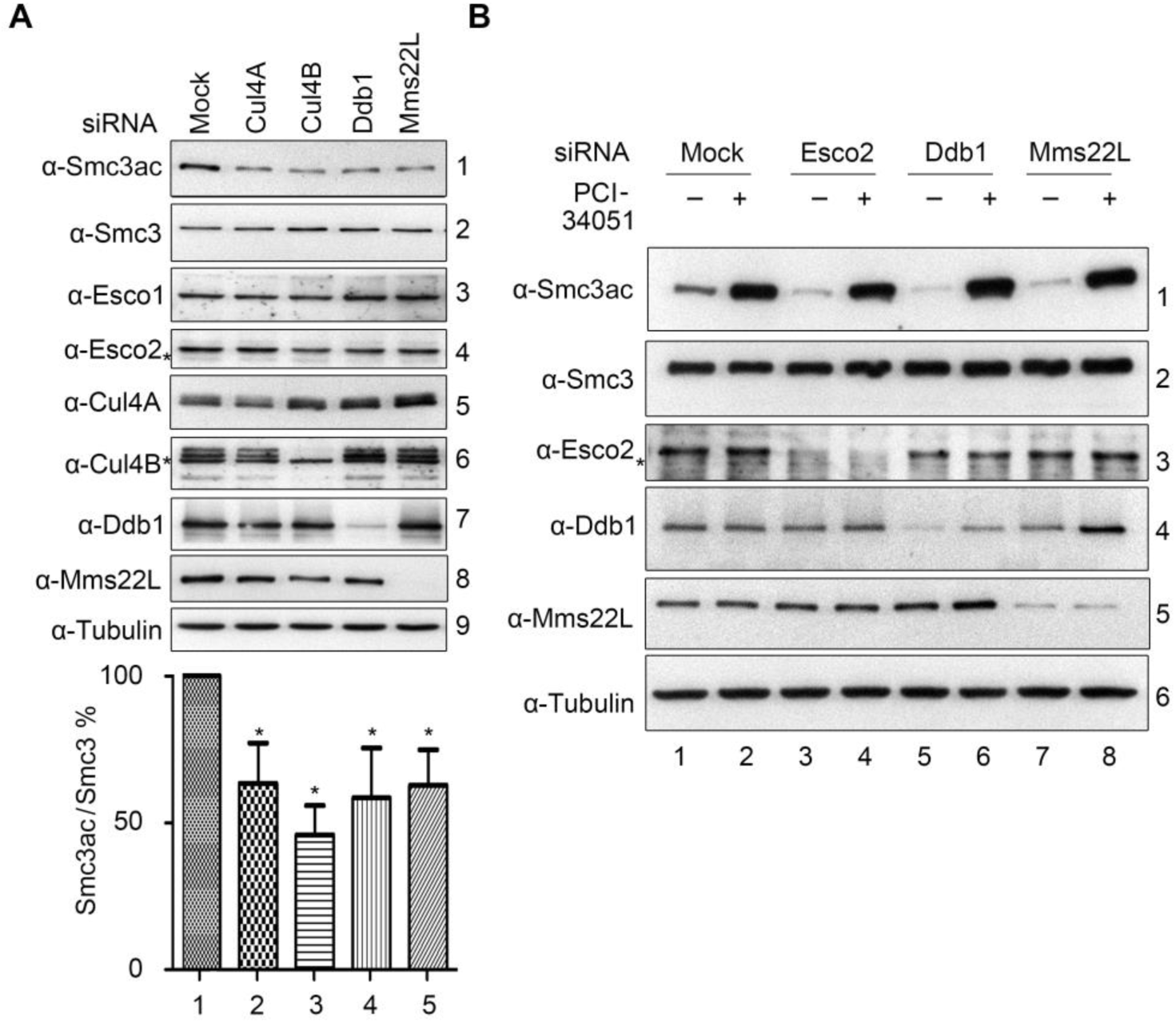
Cul4A/Cul4B-Ddb1-Mms22L are required for Esco2-dependent Smc3 acetylation. (A) Knockdown of Cul4-Ddb1-Mms22L leads to reduced Smc3 acetylation. Smc3 acetylation was analyzed in the siRNA-transfected cells as above. The statistical significance from at least three independent repeats was calculated via student’s *t*-test, *P<0.05. The results of at least three independent biological experiments were summarized in the histogram. See S4A Fig for the linear ranges of quantitation analysis of immunoblots, S4B Fig for the dosage suppression results of Esco2-depleted cells, and S4C-S4E for biological repeats. (B) Inhibition of deacetylase HDAC8 restores Smc3ac levels in Ddb1 and Mms22L – depleted cells. The indicated cells were cultured and treated by PCI-34051 for 2 h before collection and Giemsa analysis. Mitotic cells with normal cohesion were counted as described in Fig 1. The statistical significance was calculated via student’s *t*-test, ** P<0.01.

To further validate the role of CRL4s in Smc3 acetylation, we then asked whether inhibition of HDAC8 is able to restore compromised Smc3 acetylation caused by CRL4^Mms22L^-depletion. Since HDAC8 is the deacetylase of Smc3 [39], we treated proliferating cells with the HDAC8 inhibitor, PCI-34051. In the presence of PCI-34051, Smc3 acetylation increased markedly in both WT and Esco2-depleted cells (Fig 5B, lanes 1-4), as reported previously [32]. There is a similar increase in the Smc3ac level when Ddb1 and Mms22L were depleted individually (lanes 5-8). Nevertheless, Shirahige’s group shows that PCI-34051 is not able to restore the Smc3ac levels caused by compromised Pds5A-Pds5B-Esco1 branch [32]. This supports that Ddb1 and Mms22L function in the Esco2-catalyzed Smc3 acetylation pathway, which can be reversed by HDAC8. Taken together, these data reinforce the notion that CRL4^Mms22L^ ligases modulate the activity of Esco2 on Smc3 acetylation.

### Both CRL4s and PCNA help to stabilize Esco2 on chromatin

Next, we directly tested whether the regulation of CRL4s on the Esco2 activity depends on their interactions shown in Fig 3. To this end, we constructed several *Esco2* alleles defective in either CRL4s-binding (*Esco2*-LG) or PCNA-binding (*Esco2*-PIP) according to highly conserved sites from yeast to human (Fig 6A). To obtain a catalytic-deficient enzyme, we also introduced the missense mutation W539G in Esco2, which occurs frequently in Roberts Syndrome (RBS) patients [40]. Indeed, the W539G allele showed a substantially decrease in Smc3 acetylation (Fig 6B, lane 5), in agreement with its location within the acetyltransferase domain. *Esco2*-LG exhibited a significant decrease in Smc3 acetylation and cohesion efficacy to a similar extent as the catalytic-deficient W539 allele (Fig 6B, lane 3), indicating that CRL4s-mediated interaction is crucial for Esco2 operating on Smc3. Similarly, a PCNA-interaction defective allele, *Esco2*-PIP, reduced Smc3 acetylation and cohesion efficacy as well (lane 2). Interestingly, when we combined both mutations on Esco2 (Esco2-LG-PIP), we found a synergistic loss of Smc3 acetylation and cohesion (Fig 6B, lane 4). These data suggest that the function of Esco2 is cooperatively regulated through its dual interaction with both CRL4^Mms22L^ and PCNA.

**Fig 6.**
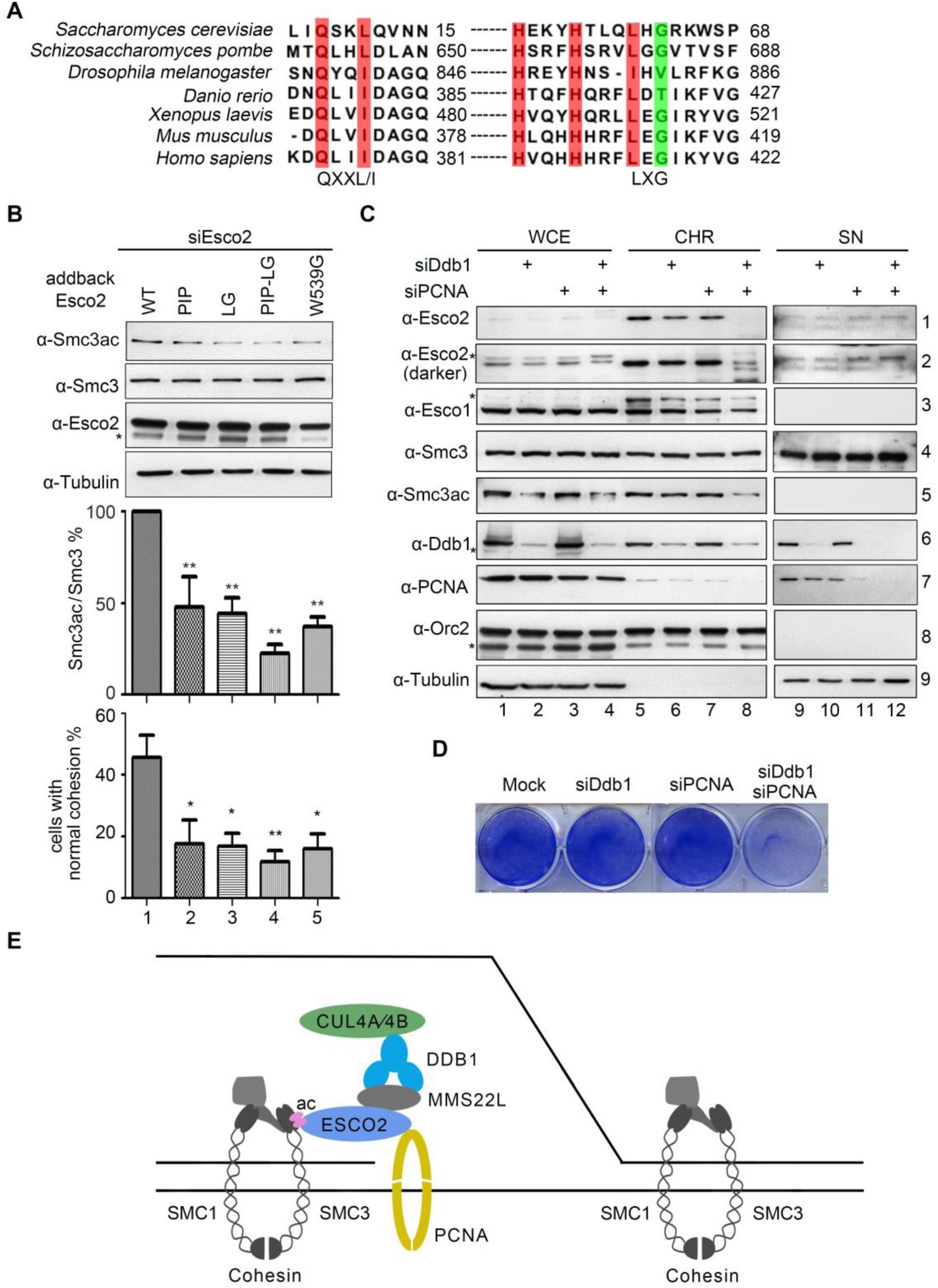
Both CRL4s and PCNA mediated interactions are required for stabilizing Esco2 on chromatin. (A) The interaction motifs of Eco1/Esco2 with PCNA and CRL4s are evolutionarily conserved from yeast to human. The amino acid sequences of Eco1/Esco2 from the indicated organisms were aligned by CLC Genomics Workbench 3. (B) *Esco2* mutants display compromised Smc3 acetylation and thereby cohesion establishment. *Esco2* alleles: PIP, *Esco2*-Q374AI376AI377A; LG, *Esco2*-L415DG417D; PIP-LG, *Esco2*-Q374AI376AI377A-L415DG417D. Quantitation from three biological repeats is shown for Smc3ac (middle) and cohesion efficiency (lower). The statistical significance was calculated via student’s *t*-test, *P<0.05,**P<0.01. The results of at least three independent biological experiments were summarized in the histogram. (C) Ddb1 and PCNA are required for the chromatin association of Esco2. Chromatin fractions were prepared from cells transfected with the Ddb1 or PCNA siRNAs. Orc2 served as a loading control of the chromatin-enriched fraction (CHR). Three independent biological experiments were performed and one of the results was shown. (D) Combined depletion of Ddb1 and PCNA caused cell death. Cells were stained with Coomassie brilliant blue 48 h after transfection with the indicated siRNAs. Three independent biological experiments were performed and one of the results was shown. (E) A co-regulation model of Esco2 by CRL4s and PCNA in human cells. As replication fork proceeding during S phase, Esco2 is recruited via interactions with fork components including PCNA and CRL4^Mms22L^ ligases. This is required for efficient Smc3 acetylation of the pre-loaded cohesin ring, which triggers the establishment of cohesion between two newly synthesized sister chromatids.

Since both CRL4^Mms22L^ and PCNA associate with replication forks, we then analyzed the contribution of Ddb1 and PCNA to the chromatin recruitment of Esco2. Chromatin fractions were prepared from the 293T cells transfected with siRNAs specific to Ddb1 or PCNA. Depletion of either Ddb1 or PCNA led to moderately reduced amounts of Esco2 on chromatin (Fig 6C, lanes 6 and 7). The combinational depletion of Ddb1 and PCNA only posed a subtle effect on the total Esco2 levels (Fig 6C, WCE, lane 4). Nevertheless, a clear synergistic loss of Esco2 on chromatin was observed (CHR, lane 8). Consistently, the level of acetylated Smc3 largely reduced whereas the total Smc3 protein on chromatin remained nearly unaffected (Fig 6C, descending panels 3 and 4). Meanwhile, the chromatin-associated Esco1 level only displayed a mild change (Fig 6C, lane 8, panel 2). In good agreement with this, only when Ddb1 and PCNA depletions were combined, dramatic cell death was observed by live cell staining (Fig 6D). These data suggest a cooperative mechanism for CRL4s and PCNA to properly target the essential cohesin acetyltransferase Esco2 on its substrate Smc3, which contributes to the coupling between the establishment of sister chromatid cohesion and replication fork progression in human cells (Fig 6E).

## Discussion

How sister chromatid cohesion is established in mammals remains largely unclear. Here we have identified an evolutionarily conserved mechanism of CRL4 ubiquitin ligases, together with PCNA, in regulation of DNA replication-coupled cohesion establishment in human cells.

The essential step to establish cohesion is Smc3 acetylation by Eco1 and Esco in yeast and human, respectively [13–15]. Precise control of the reaction is required for this essential cellular process. One of the main findings of this study is that human CRL4^Mms22L^ ligases exclusively interact with and preferentially regulate Esco2. Despite that both Esco1 and Esco2 catalyze acetylation of Smc3, their temporal regulation is distinct [32, 37, 41]. Esco1 acetylates Smc3 in a Pds5-dependent manner before and after DNA replication [37], whereas Esco2 is believed to function during S phase. Our findings provide molecular details of how Esco2 is controlled in a DNA replication-coupled fashion by dual interaction with CRL4^Mms22L^ and PCNA in human cells.

During the revision of this manuscript, Peters’s group reported that Esco2 is recruited to chromatin via direct association with MCM, the core of eukaryotic replicative helicase Cdc45-Mcm2-7-GINS [42]. It’s worth noting that the contributions of MCM, PCNA and CRL4s to Esco2 regulation are not mutually exclusive. A very interesting finding in their work is that Esco2 binds MCM predominantly in the context of chromatin regardless that there are largely excess amounts of MCM in nucleoplasm [42]. Even among the abundant chromatin-loaded MCM rings, only a small portion of these are activated and assembled into replication forks [43, 44]. How Esco2 is specifically recognized by the activated MCM and travels with replication fork has therefore not been addressed yet. The interactions of Esco2 with PCNA and CRL4^Mms22L^ identified previously and in this study [19], albeit relatively weak or transient, may contribute to the preferential association of Esco2 with the activated MCMs on replication fork. Intriguingly, in HeLa cells, Mms22L-TONSL bind MCM as well as replication-coupled H3.1-H4 [45–50]. Therefore, it will be of great interest to test the functional interplay among these fork-associated factors in future.

In addition to these interactions, CRL4s have been found to be involved in multiple replication-coupled chromatid events. For instance, Cul4-Ddb1 (Rtt101-Mms1) ubiquitylates histone H3-H4, which elicits the new histone hand-off from Asf1 to other chaperones for chromatin reassembly in both yeast and human cells [34]. Another CRL4, CRL4^WDR23^, ubiquitylates SLBP to activate histone mRNA processing and expression during DNA replication [51]. Further studies are needed to illustrate the details of crosstalk among these replication-coupled events, for instance, nucleosome assembly and cohesion establishment÷ in human cells.

Over-expression of Cul4, Ddb1 and Mms22L has been reported to correlate with lung and esophageal carcinogenesis [52], implicating them as key genome caretakers. Mutations in *Esco2* gene cause Roberts Syndromes with a predisposition to cancer [40]. The functional interplay between Cul4-Ddb1 and Esco2 identified here will shed new light for us to understand the etiology of these human diseases.

## Materials and methods

### Cell culture and RNAi

HEK293T cells were cultured in DMEM media supplemented with 10% fetal bovine serum (FBS, Gibco) at 37°C with 5% CO_2_. For RNAi experiments, cells were transfected with 80 nM siRNAs using Lipofectamine 3000 (Invitrogen) for 48 h following the manufacturer’s instructions. Over-expression plasmid or control plasmid for target genes was transferred into the cells when necessary. For HDAC8 inhibition experiments, 6.25 μM PCI-34051 (Selleckchem) was applied 3 h before harvest. Immunoblotting with specific antibodies was used to confirm the downregulation of the targets. The sequences of siRNA oligos used in this study are listed in Table 1. All siRNA oligos were synthesized by Sangon Biotech, China.

**Table 1:**
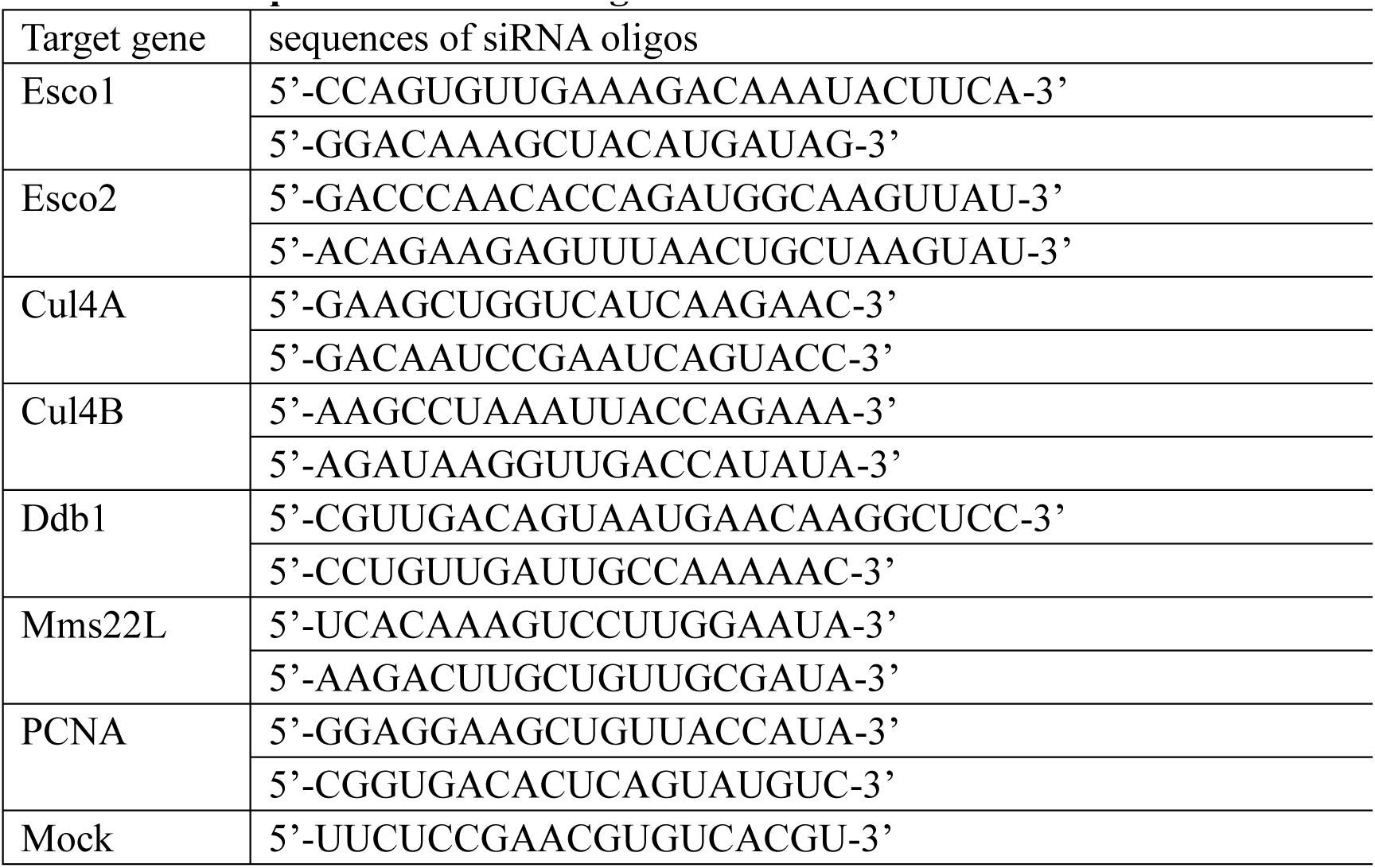
The sequences of siRNA oligos.

### Plasmid construction

Trizol reagent (CWBIO) was used to isolate total RNA according to the manufacturer’s instructions. cDNA was synthesized using reverse transcriptase (Promega). Full length genes studied were inserted to the prk5-Flag, GFP or mCherry vectors [53]. The prk5-Flag vector was kindly provided by Dr. Jun Tang (China Agricultural University) and Flag was replaced by GFP or mCherry when necessary.

### Chromosome spreads

Chromosome spreads were performed as described in [54], with minor modifications. In brief, cultured cells were harvested by trypsinization and then 75 mM KCl was used as hypotonic treatment. Cells were fixed with methanol and acetic acid (3:1) three times and then dropped onto the slides. After half an hour, cells were stained with 0.05% Giemsa (Merck) for 10 min at room temperature. Images were captured using a Leica microscope equipped with a 100×/NA1.3 oil objective. The incidence of sister chromatid separation was determined from at least 200 mitotic cells and all experiments were repeated at least three times. In our experimental conditions, almost all chromosomes within a single cell display similar cohesion defects. Total cohesion defects were counted as cells exhibiting precocious separation of both arms and centromeres, while *CEN* cohesion defects shown in the Supporting Figures were calculated for the percentages of cells bearing separated centromeres among mitotic cells.

### Cell extract, Immunoprecipitation and Immunoblotting

Cells were washed twice with PBS. To obtain whole cell extracts for immunoblotting, cells were resuspended with RIPA buffer (50 mM Tris-HCl, 250 mM NaCl, 1% TritonX-100, 0.25% Sodium deoxycholate, 0.05% SDS, 1 mM DTT) and lysed on ice for 20 min. For immunoprecipitation experiments, cells were resuspended in lysis buffer (50 mM Tris-HCl, 150 mM NaCl, 1% NP-40, 5 mM EDTA, 10% glycerin) and incubated on ice for 20 min, then sonicated for 30 sec. For each sample, 250 μg total protein was incubated with anti-Flag agarose for 1.5 h at 4°C, then washed five times with lysis buffer. All of the samples were run on sodium dodecyl sulfate polyacrylamide gel electrophoresis (SDS-PAGE) and transferred to PVDF membranes. Signals were detected with specific antibodies using eECL Western Blot Kit (CWBIO).

### Affinity purification coupled to mass spectrometry (AP-MS)

Esco2-HF was purified from whole cell extracts by anti-Flag M2 (Sigma) and Ni^2+^ affinity gels successively. Nonspecific bound proteins were removed by washing with 0.25 μg/μl Flag peptide. Bound fraction was eluted by 2 μg/μl Flag peptide and 300 mM imidazole, respectively. An untagged cell line and Esco1-HF were subjected to the same procedure as controls. The final eluates were analyzed by mass spectrometry analysis (Q Exactive™ Hybrid Quadrupole-Orbitrap Mass Spectrometer, Thermo Fisher). The procedures were repeated three times for both Esco2-HF and Esco1-HF to identify the different interactors of Esco2 and Esco1.

### Fluorescence stain

Cells were grown in cover glasses placed into 6-well plates (Nunc) and transfected with plasmids using Lipofectamine 3000 (Invitrogen) for 24 h according to the instructions. After being fixed with 4% paraformaldehyde, cells were stained with 1 μg/ml DAPI for 5 min at room temperature. Images were captured with a laser-confocal microscope (DMi8; Leica Microsystems).

### Antibodies

Antibodies used in this work were as below: Esco1 (Abcam, ab180100), Esco2 (Abcam, ab86003), Cul4A (proteintech, 14851-1-AP), Cul4B (Proteintech, 12916-1-AP), Ddb1 (Abcam, ab9194), Mms22L (Abcam, ab181047), Smc3 (BETHYL, A300-060A), acetylated Smc3 (Merck, MABE1073), Orc2 (CST, #4736), Tubulin (MBL, PM054) and PCNA (Santa Cruz, sc-56).

## Acknowledgments

We thank Dr. Judith L. Campbell for improving the manuscript, Drs. Cong Liu, Jun Tang, Qun He and members of the Lou lab for discussion. We are also grateful to the anonymous reviewers for constructive suggestions.

## Author Contributions

H.L., J.J.Z. and Q.C. conceived and designed the overall project; H.S. conducted most experiments with the help from J.X.Z. and J.J.Z.; Z.L. helped in mass spectra analysis; H.L. H.S. and Q.C. wrote the manuscript with input and editing from all of the authors.

